# Subsequent memory effect in the inferior frontal gyrus revealed by fNIRS

**DOI:** 10.1101/2025.10.02.679993

**Authors:** Petra Bíró, Silvy HP Collin

**Affiliations:** Department of Computational Cognitive Science, Tilburg School of Humanities and Digital Sciences, Tilburg University, Tilburg, the Netherlands

## Abstract

A central finding in memory research is the subsequent memory effect, which describes consistent neural differences during encoding between events that are later remembered versus those that are forgotten, which has been reliably replicated with both EEG and fMRI for the past decades. Replicating the subsequent memory effect using fNIRS could enable research opportunities that are difficult to pursue with other methods, including studies with children, patient populations, or experiments in highly naturalistic settings. Therefore, our study investigated whether the prefrontal cortex is differentially involved in subsequently remembered versus subsequently forgotten stimuli using fNIRS. Our results showed that in particular channels mapping onto the inferior frontal gyrus showed more activation during encoding of subsequently remembered events compared to subsequently forgotten events. These results demonstrate that fNIRS can reliably capture the subsequent memory effect, providing new opportunities to study memory mechanisms across diverse populations and real-world contexts.

## 1. Introduction

Understanding why some experiences are remembered while others are forgotten is a central question in cognitive neuroscience (Sanquist et al., 1980; Paller et al., 1987). The subsequent memory effect is one of the core findings in memory research, and refers to the reliable neural differences observed at encoding between events that are later remembered and those that are later forgotten (Mecklinger and Kamp, 2023; Kim, 2019; Maillet and Rajah, 2014; Kim, 2011). It has been robustly replicated across methodologies; EEG (Li et al., 2024; Wiemer et al., 2021; Kim et al., 2020; Piñeyro Salvidegoitia et al., 2019; Long et al., 2014; Hanslmayr and Staudigl, 2014), fMRI (Yu et al., 2025; Halpern et al., 2023; Soch et al., 2022; Wing et al., 2018; Fletcher et al., 2003; Rypma et al., 2002), and combined EEG-fMRI (Hoppstädter et al., 2015; Hanslmayr et al., 2011). Building on this foundation, our project investigates the feasibility of using a relatively new neuroimaging method –functional Near-Infrared Spectroscopy (fNIRS)– to reliably detect such a subsequent memory effect in the human brain. fNIRS is a tool that measures hemodynamic response and mostly differs from other brain activity measurement techniques in its portability, making it particularly attractive for cognitive neuroscience studies aiming to explore real-world functioning. fNIRS has been used in studies involving healthy adults and clinical patients with neurological disorders (Pinti et al., 2020). Here, our overarching goal is to measure the prefrontal cortex (PFC) during memory encoding using fNIRS equipment and relate its activity to behavioral data from a subsequent memory test. Translating the subsequent memory effect—robustly demonstrated with other neuroimaging methods (Yu et al., 2025; Li et al., 2024; Halpern et al., 2023; Soch et al., 2022; Wiemer et al., 2021; Kim et al., 2020; Piñeyro Salvidegoitia et al., 2019; Wing et al., 2018; Hoppstädter et al., 2015; Long et al., 2014; Hanslmayr and Staudigl, 2014; Fletcher et al., 2003; Rypma et al., 2002)—to fNIRS will open avenues for research not easily performed with other methods like fMRI (Seghier et al., 2019), such as studies in children, patient groups, or highly naturalistic paradigms.

In a subsequent memory paradigm, encoding trials are categorized as remembered or forgotten based on later memory performance. A meta-analysis (Kim, 2011) found that subsequent memory effects are most prominent in the inferior frontal cortex (IFC), and was particularly prominent for verbal material. Additionally, subsequent memory effects have been found in other subregions within the PFC, like dorsolateral PFC Rypma et al. (2002) and ventromedial PFC Guo and Yang (2023); Wing et al. (2018). Clinically, the subsequent memory effect has also been investigated in individuals from several types of patient groups that are known to (also) have memory-related symptoms, like Alzheimer’s disease patients (Soch et al., 2024), schizophrenia patients (Collier et al., 2014) and epilepsy patients (Sidhu et al., 2013). In an fNIRS study using a directed forgetting paradigm, more activity was observed during intentional forgetting than intentional remembering (Jing et al., 2022), which is a paradigm that indirectly relates to a subsequent memory paradigm.

In summary, both fMRI and EEG have been used to study the subsequent memory effect or similar paradigms, but there is a lack of substantial research directly investigating the subsequent memory effect using fNIRS. Our study will bridge this gap by investigating whether fNIRS measurements during memory encoding can elicit the difference between brain activity for later remembered and forgotten events. While various individual studies using in particular fMRI have discovered subsequent memory effects in various subregions within the PFC (e.g., Rypma et al. (2002) in dorsolateral PFC, and Wing et al. (2018) in medial PFC), we expected the strongest subsequent memory effect in the inferior frontal cortex, given that a large meta-analysis across as many as 74 neuroimaging studies (Kim, 2011) most consistently mentions that this part of the PFC is related to a subsequent memory effect.

## 2. Materials and Methods

This study is a re-analysis of the data from Bíró and Collin (2025) which focused on how the PFC is involved in representing schema-violating information. The present study addresses a distinct research question focused on the subsequent memory effect. For this purpose, we re-used the data from day 1 of this fNIRS experiment to investigate whether the PFC was involved in subsequent memory performance.

### 2.1 Participants

The dataset has 38 participants between 18 and 33 years old. Participants gave written informed consent before starting the experiment, and received study credits for their participation. Two participants were excluded due to technical issues during data acquisition. The experiment was approved by the Research Ethics and Data Management Committee of TSHD, Tilburg University.

### 2.2 Summary of task, stimuli and data preprocessing

Here, a summary of the task that participants performed on day 1 of this 2-day experiment, the stimuli used, and the data preprocessing steps taken. More details can be found in Bíró and Collin (2025).

In this experiment, participants were seated in front of a computer monitor wearing a g.tec fNIRS headcap (g.tec, n.d.) and encoded 32 individual stimuli (that on day 1 lasted 9 seconds per stimulus, with an 1 second intertrial interval). These stimuli were fictional activity descriptions, each related to either a holiday or educational activity. An example of such an activity description is: *Here, people cultivate and observe plants specifically adapted to thrive in lunar environments, considering low gravity and limited resources*. Besides reading the activity description, participants also viewed a corresponding map with the activity’s location circled. Descriptions were shown below the map with a title above, and all were created using ChatGPT and manually refined.

Participants’ memory for the stimuli was tested in a subsequent recognition memory test (self-paced) directly following encoding that included the actual stimuli, 32 lure stimuli and 32 completely new stimuli. In this memory test, participants were asked whether they recognized the description (i.e., old or new), whether they remember which map the stimulus was from (holiday or education) and whether they remember which location the stimulus was from (out of 9 possible location on a map). As memory performance measure, recognition of the activity descriptions was used (i.e., the old/new question).

fNIRS signals were recorded wirelessly using the g.Nautilus system with eight frontal cortex optodes at 250 Hz, using 760 nm and 850 nm LEDs, synchronized via Bluetooth with Simulink/Matlab. fNIRS data were preprocessed using MNE (Gramfort et al., 2013), including bandpass filtering for 0.01 – 0.1 Hz (Gramfort et al., 2024), epoching (0–8 s), downsampling to 10 Hz, and baseline correction using the rest block preceding the encoding task itself; E-prime trial sequences were exported, and added as epoch metadata for analysis in R.

Our preprocessing pipeline for both fNIRS and behavioral data was identical to (Bíró and Collin, 2025). More details concerning the stimuli, task and data preprocessing can therefore be found in the methods section of Bíró and Collin (2025).

### 2.3. Statistical analysis

To address our research question, binary logistic regression was employed (Cox, 1958) to examine whether brain activity measured through fNIRS could predict subsequent memory performance. The dependent variable in the analysis, *remembered_forgotten*, indicates whether a participant recalled a specific item or not. The independent variable, *value*, represents the continuous HbO (oxy-hemoglobin) signal recorded during encoding on day 1. A binary logistic regression model was run on the aggregate data, i.e., all channels combined. For this model, we used the data from time 4 to 6 given that this is the approximate time-window in which we expected the peak of the hemodynamic response function (HRF) related to the BOLD response. Before running the statistical analysis, we calculated the average HbO concentration across all trials in the task and excluded everyone that on average was 2 standard deviations or more away from the group mean as being outliers (which was the case for 4 participants).

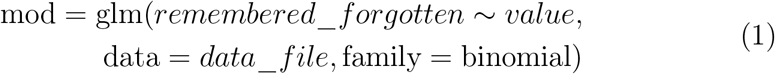

Besides this binary logistic regression on the aggregate data across all channels, separate (identical) models were run for each channel individually to explore the specific contribution of individual PFC regions to memory prediction. We expected in particular a subsequent memory effect in channels 2 and 6, given that they map mostly onto the inferior frontal gyrus (see Bíró and Collin (2025), Table 1).

## 3. Results

We investigated whether brain activity during encoding can predict subsequent memory performance. We primarily focused on the channels mapping onto the inferior frontal gyrus (i.e., channels 2 and 6), given that a meta-analysis (Kim, 2011) showed this to be the region most consistently mentioned in relation to the subsequent memory effect. Channel-wise data showed that channel 6 –mapping onto the left inferior frontal gyrus– had significantly higher activity in subsequently remembered compared to subsequently forgotten trials (b = 0.038, SE = 0.016, z = 2.384, P = 0.017, see Figure 1). The right inferior frontal gyrus (i.e., channel 2) did not reach significance (b = 0.006, SE = 0.01, z = 0.635, P = 0.526). Thus, it was in particular the left inferior frontal gyrus that showed a subsequent memory effect in our dataset (i.e., P *<* 0.025, corrected for multiple comparisons due to separate analyses for left and right hemispheres).

**Figure 1:**
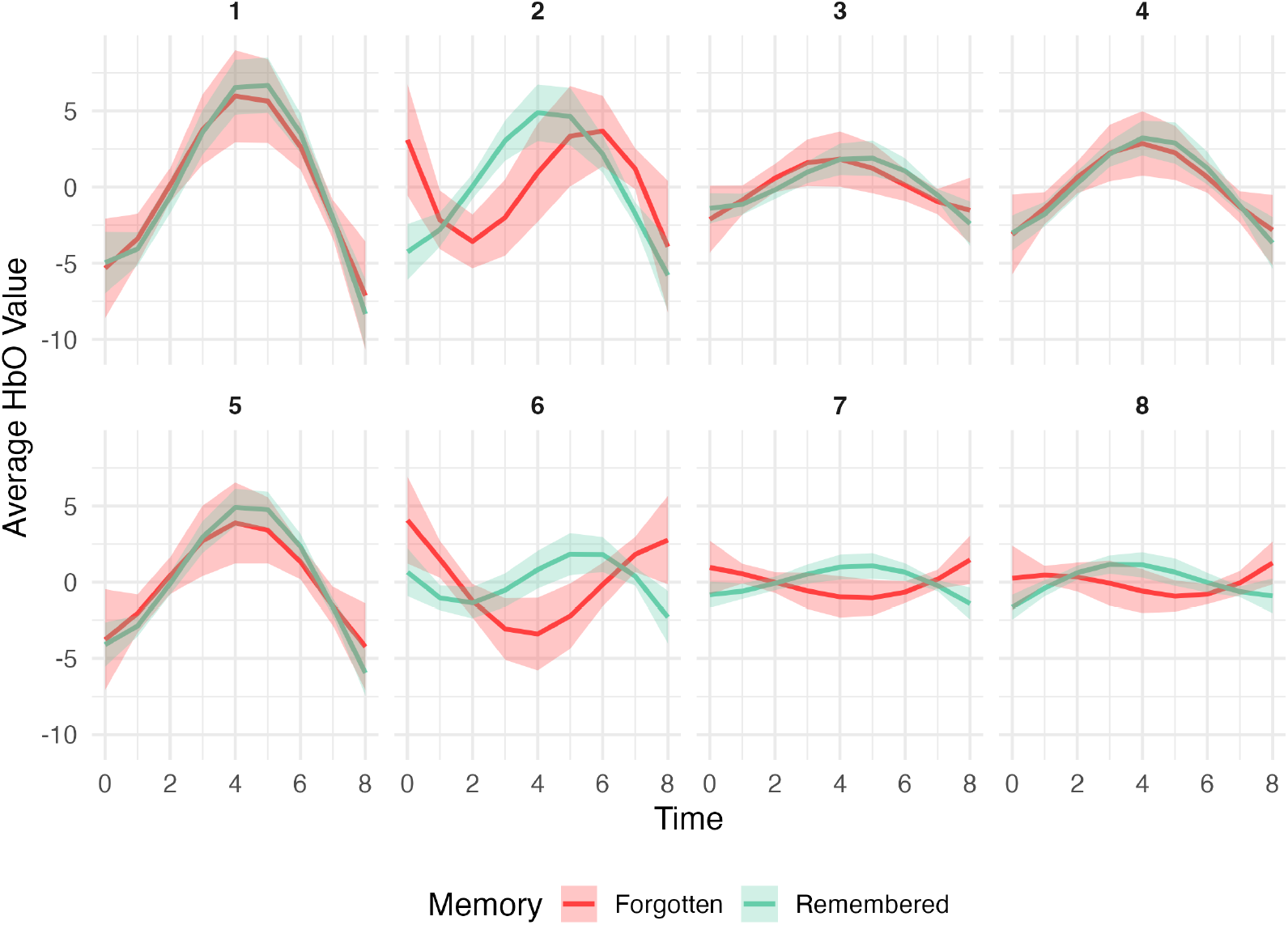
Channel-wise HbO levels for remembered (green) and forgotten (red) trials (mean ± S.E.M.). Time-point 0 is the onset of the stimulus. Channels 1 and 5 map onto the medial PFC, channels 2 and 6 onto the inferior frontal gyrus, and channels 3, 4, 7 and 8 onto the dorsolateral PFC (see Bíró and Collin (2025), Table 1).

Figure 2 shows the aggregate data where the HbO concentration levels are averaged across all 8 channels. This shows that the HbO concentration levels across the entire PFC for trials that were subsequently remembered on day 1 are numerically higher than for subsequently forgotten trials, but this difference across the entire PFC did not reach significance (b = 0.026, SE = 0.019, z = 1.31, P = 0.19). These results suggest that it is primarily the (left) inferior frontal gyrus that is implicated in the subsequent memory effect in this fNIRS dataset.

**Figure 2:**
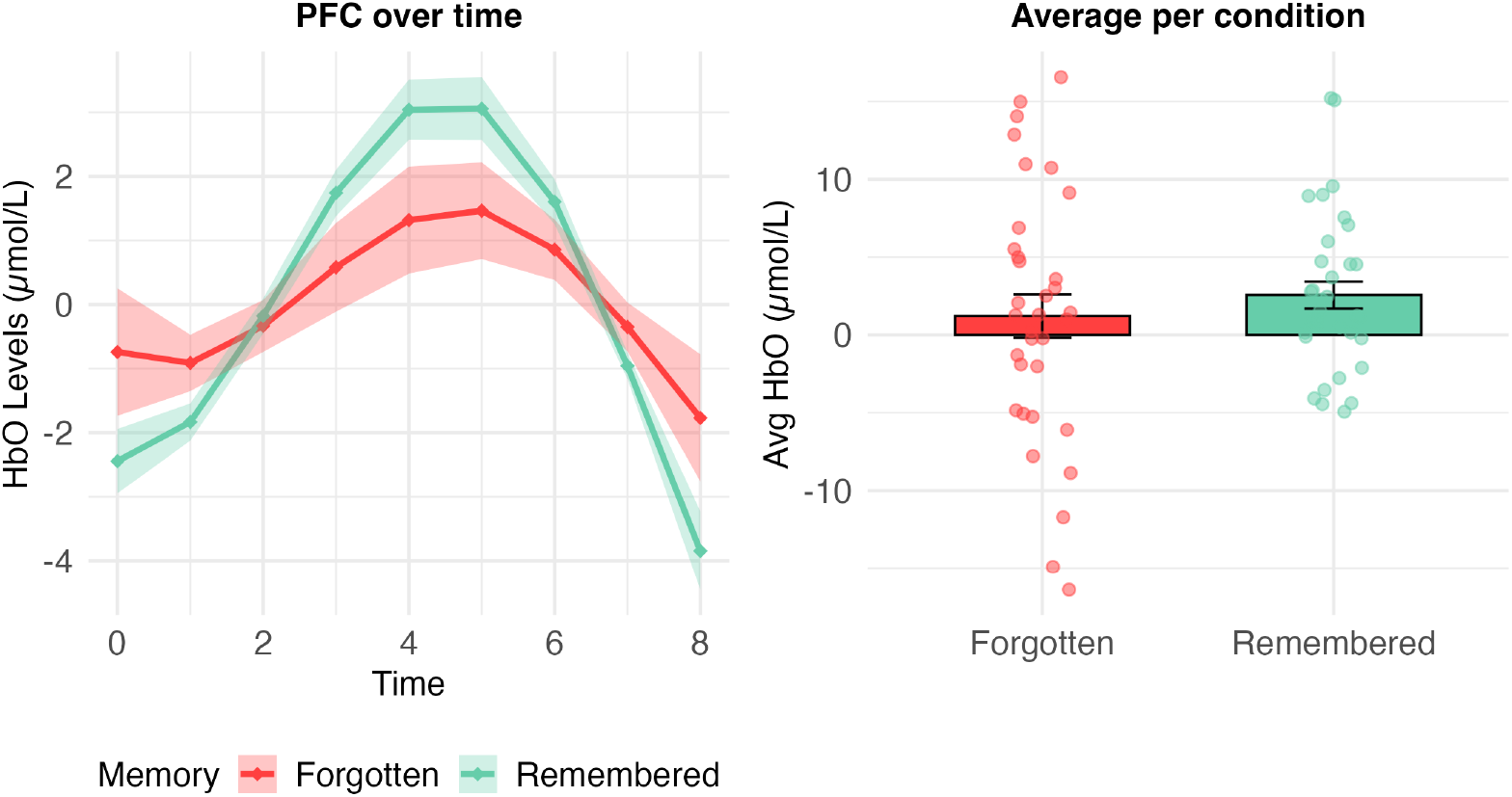
HbO concentration for remembered (green) and forgotten (red) trials, averaged across trials and channels. Left: HbO concentration timeline (mean ± S.E.M.). Time-point 0 is the onset of the stimulus. Right: A bar plot representing the mean ± S.E.M. overlaid with data from individual participants, averaging time-points 4, 5 and 6 (being the expected peak of the HRF BOLD response).

## 4. Discussion

Here, we set out to replicate the subsequent memory effect (as discovered using fMRI) using low-density, portable fNIRS equipment. Our results revealed that it is possible to reliably detect a subsequent memory effect in the left IFG using low-density fNIRS. This suggests that the subsequent memory effect is a robust measurement that is detectable across multiple modalities. The fact that fNIRS is able to reliably detect the subsequent memory effect is useful for naturalistic experiments that would need to rely on a mobile experimental set-up.

These results are in line with IFG being considered one of the key brain regions involved in subsequent memory effects (Kim, 2011). However, it is in contrast to fMRI research that also showed a subsequent memory effect in dorsolateral PFC (which would map onto channels 3, 4, 7 and 8) (Rypma and D’Esposito, 2003) and ventromedial PFC (which would map onto channels 1 and 5) (Guo and Yang, 2023; Wing et al., 2018). One possible explanation for this discrepancy is that low-density fNIRS has limited coverage and depth sensitivity, making it less suited to detect more distributed or deeper prefrontal contributions. It is likely that our optodes only covered part of the dorsolateral and medial PFC, which means that it is harder to reliably draw conclusions from these regions using low-density fNIRS.

An important contribution of the present study is methodological: our findings demonstrate that the subsequent memory effect can be detected with a low-density fNIRS setup. While such systems offer limited spatial resolution compared to fMRI, they have clear advantages in terms of cost, portability, and tolerance for movement. The fact that a reliable subsequent memory effect was observed under these suboptimal conditions suggests that fNIRS can be a practical alternative for studying memory processes in contexts where fMRI is not feasible, such as with infants, older adults, or in naturalistic and field settings.

### 4.1. Limitations and future directions

Our results only showed evidence for a subsequent memory effect in the left IFG, and not in the right IFG. Kim (2011) suggests that there is some evidence that encoding of verbal material is more focused on left hemisphere involvement in the context of subsequent memory analysis, while pictorial stimuli activate more regions in the right hemisphere. Since this experiment included both verbal information (i.e., the event descriptions that the participant read) and pictorial information (i.e., the map with an indication of the specific location), it is difficult to determine whether participants on average focused more on the image or the text. The fNIRS results suggest that participants might be focusing more on the text, but to make firm conclusions about this lateralization, future research could use an experimental design where verbal and pictorial information are clearly differentiated in time.

Our results suggest that fNIRS may serve as a viable tool for investigating memory processes. As indicated above, this would be especially relevant in situations where fMRI is impractical, such as with infants, older adults, or in more naturalistic and field environments. Our study, however, only tested healthy young adults. Thus, future research is necessary to translate our subsequent memory effect findings to such naturalistic environments and/or to children and older adults.

## 5. Conclusion

Overall, our findings support the feasibility of using low-density fNIRS to study the subsequent memory effect, opening possibilities for investigating memory processes in populations and environments where traditional neuroimaging is not practical.

## Author CRediT statement

**Petra Biro**: Conceptualization, Software, Data Curation, Investigation, Validation, Formal analysis, Visualization, Writing - Original Draft. **Silvy H.P. Collin**: Conceptualization, Formal analysis, Visualization, Supervision, Project administration, Writing - Review and Editing.

## Acknowledgements

We thank members of the DAF Technology lab at which we ran data acquisition of this project. In particular, we thank Hans van den Dool for help regarding IT and fNIRS hardware/software set-up in the fNIRS lab.

## Notes

### Competing Interest Statement

The authors have declared no competing interest.

